# Programmable microactuators phase-lock cilia to local oscillatory flow

**DOI:** 10.64898/2026.05.14.725102

**Authors:** Farzin Akbar, Veikko F. Geyer, Benjamin M. Friedrich, Maximilian Kotz, Stefan Diez, Mariana Medina-Sánchez

**Affiliations:** B CUBE – Center for Molecular Bioengineering, TUD Dresden University of Technology (Dresden-Germany); Cluster of Excellence Physics of Life, TUD Dresden University of Technology (Dresden, Germany); Max Planck Institute of Molecular Cell Biology and Genetics (Dresden, Germany); CIC nanoGUNE BRTA (San Sebastián, Spain); Ikerbasque, Basque Foundation for Science (Bilbao, Spain)

## Abstract

Hydrodynamic synchronization of motile cilia is essential for biological functions such as fluid transport, locomotion, and developmental patterning. It comprises the generation and the response to local flows in complex geometries. Besides their central role in physiology, direct experimental tests of ciliary responses to local flows at cellular length and time scales have remained elusive, largely due to the absence of tools capable of applying controlled, and localized flow stimuli. Here, we introduce programmable, nanometer-thin Ti/Pt microactuators that generate well-defined hydrodynamic forcing at biologically relevant frequencies while operating at biocompatible sub-Volt voltages. This platform is pioneering a controlled local hydrodynamic stimulation of individual motile cilia. We quantify the flow fields and forces produced by single microactuators using particle image velocimetry. Applying local oscillatory flows close to motile cilia of the green alga Chlamydomonas reinhardtii, we probe their dynamic response by quantifying phase-locking between cilia and microactuators. This quantification is aided by combining machine-learning–based image segmentation, oscillator phase reconstruction, and circular statistics. During actuation, we observe signatures of phase-locking: those include a reversible modulation of the fluctuations in phase-difference between cilium and actuator and a systematic shift in ciliary beating frequency. Beyond providing a bio-compatible and precise platform for local hydrodynamic stimulation, our approach establishes an experimental framework for directly testing theories of hydrodynamic synchronization and load adaptation in systems of motile cilia.

## 1 INTRODUCTION

Motile cilia are hair-like cell appendages that beat periodically to move fluid or propel motile cells in a liquid (*1*). On airway epithelia and numerous other tissues, arrays with thousands of cilia synchronize their beat into metachronal waves with fixed phase relationships between nearby cilia (*2–4*) suggested to enhance fluid transport and mixing (*5,6*). Direct hydrodynamic interactions between cilia were proposed as potential synchronization mechanism (*7*), which was experimentally confirmed recently using pairs of ciliated cells held at a distance (*8*). Computer simulations indeed showed that hydrodynamic interactions are sufficient to drive metachronal coordination in cilia arrays (*5, 9–12*). Yet, if hydrodynamic interactions are the only synchronization mechanism remains debated (*13–15*). This motivates controlled, well defined flow perturbation experiments. The bi-ciliate single-cell green alga *Chlamydomonas reinhardtii* became a premier model to study synchronization (*13, 16–20*). Exposing *Chlamydomonas* cells to global oscillatory flow in microfluidic devices showed that cilia phase-lock their beat to the external flow (*13, 20*) (later confirmed for cilia bundles (*21*)). Yet, biological variability and active cilia phase noise (*22*) perturb synchronization (*17, 23*), resulting in phase-slips in pairs of synchronized cilia (*17, 18*) and locally synchronized domains in cilia carpets (*3*). Thus, dissecting the *physical* mechanisms under physiological conditions remains challenging. In addition, key control parameters such as local forcing strength, detuning, waveform, and geometry (distance/orientation relative to the cilium) are difficult to modify in biological cells, and global-flow setups provide limited spatial selectivity. On the other hand, microactuators (including artificial cilia) can mimic ciliary strokes. To this end, a variety of microscale actuation strategies have been demonstrated, including pressure-driven (*24*), magnetic (*25,26*), electrostatic (*27*), optical (*28*), and electrochemical actuation (*29–33*). Taken together, microactuators provide a complimentary approach for imposing controlled *in situ* perturbations on motile cilia.

Recent results highlight the promise of these strategies for lab-on-chip fluid transport, active-matter studies, and biomimicry investigations (*34, 35*). Among these, ultrathin surface-electrochemical actuators based on platinum films as bilayer (e.g., Ti/Pt) are particularly attractive (*33*). Surface oxidation/reduction reactions generate differential surface stresses that bend the microactuator at low voltages in aqueous media, enabling microscale actuation at biologically relevant frequencies. Their compatibility with microfabrication techniques enables them as a scalable platform for biohybrid hydrodynamic studies (*33*). Here, we introduce a biohybrid platform that couples cilia of living *Chlamydomonas reinhardtii* cells to programmable Ti/Pt microactuators operating at biologically relevant frequencies, to probe frequency entrainment and hydrodynamic phase-locking under controlled conditions. **FIGURE 1A** displays the experimental setup, where an inverted optical microscope with phase-contrast is equipped with a microinjector and a micromanipulator. The microinjector provides finely controllable suction and pressure to gently aspirate a single cell onto the tip of a glass microcapillary and hold it there without damage. The three-axis micromanipulator then enables precise positioning of the microcapillary and therefore the attached cell in *x, y*, and *z*, allowing accurate placement relative to the microactuator. A source-measurement unit (SMU) is programmed to deliver voltage square waveforms to operate the microactuators. A high-speed camera is used to record movies of microactuators and cells (see Materials and Methods). **FIGURE 1B** displays a custom 3D-printed chamber, in which the working Pt electrode, counter Pt electrode, and Ag/AgCl reference electrode is positioned together with the microcapillary used to hold the *Chlamydomonas* cell next to the microactuator. **FIGURE 1C** displays a close-up of the setup, with the working principle of the Ti/Pt bilayer microactuator shown as inset: When the microactuator is in an electrolyte solution and its potential is raised to about +1 V *versus* an Ag/AgCl reference, the exposed Pt surface is oxidized to PtOx. This oxidation inserts oxygen species into the platinum surface and effectively makes the thin Pt film swell. Thus, during electrochemical oxidation and reduction, the platinum layer expands and contracts, respectively. Because the Pt is only on one side and the opposite side is passivated with a Ti layer, this expansion results in a flattening deformation of the whole strip. When the potential is driven again down to about −0.2 V *versus* Ag/AgCl, the PtOx is reduced back to Pt, hence the Pt layer contracts, and the microactuator attains again its initial bent state corresponding to this voltage.

**FIGURE 1:**
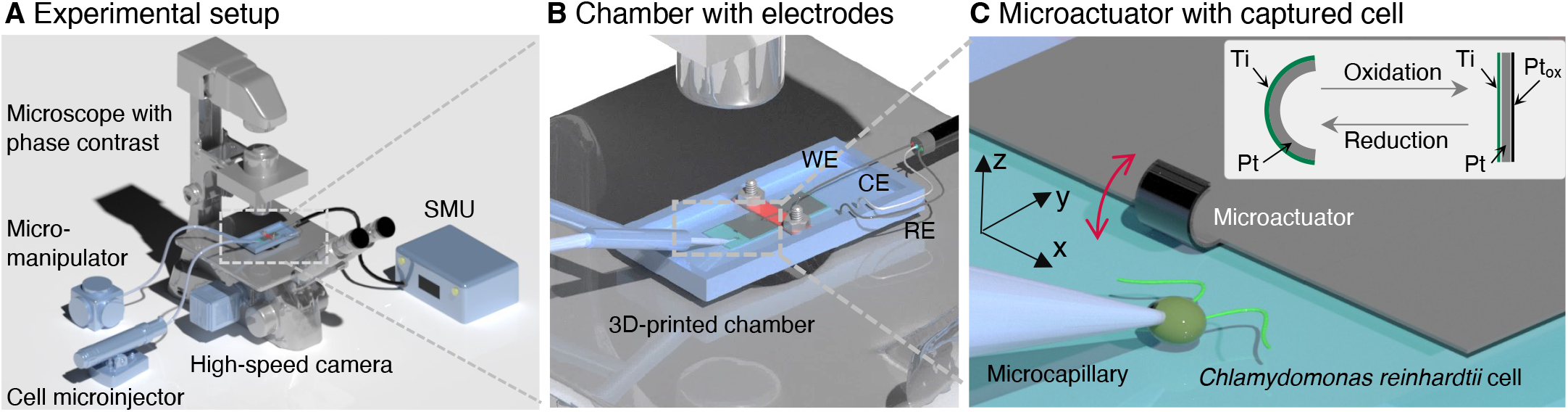
Schematic of the experimental setup used to study electrochemically driven actuation near single cells. **(A)** Overview of the inverted phase-contrast optical microscope equipped with a microinjector to hold the cell, a micromanipulator to position the cell, a high-speed camera, a source-measurement unit (SMU), and a 3D-printed chamber mounted on the microscope stage. **(B)** Magnified view of the chamber showing the working (WE), counter (CE) and reference (RE) electrodes for electrochemical stimulation. **(C)** Close-up of the microcapillary positioning a single living *Chlamydomonas reinhardtii* cell in front of the oscillating microactuator. Cells were held on the tip of a microcapillary where suction was generated by a microinjector. Inset: schematic of the Ti/Pt bilayer microactuator, where reversible oxidation and reduction of platinum drive the bending of the microactuator.

The Ti/Pt microactuators were produced on 0.5 mm-thick 50 mm × 50 mm borosilicate glass wafers, with 18 chips on each glass wafer and hundreds of actuators on each chip, using standard photolithography and thin-film deposition. (Error! Reference source not found.**AB**). The active bilayer consists of an 18 nm Ti/Pt stack, optimized for its large bending amplitude, which is released by etching an underlying aluminum sacrificial layer (Error! Reference source not found.**C)**. In a three-electrode configuration driven by custom waveforms, these microactuators are operated in Tris–acetate–phosphate (TAP) medium (*36*) at various frequencies with sub-Volt amplitudes, allowing flows to be switched on and off and the drive frequency to be controlled precisely. Actuator-generated flows are quantified by tracking particles inside the surrounding electrolyte, and, after flow characterization, individual *Chlamydomonas* cells are positioned near the microactuator and imaged at high speed (230 frames per second). From these recordings, cilia beating in the presence and absence of microactuator-generated local flows is quantified and phase-locking is demonstrated.

## 2 Results

To characterize the flow field generated by one microactuator, we added 0.5-μm-diameter polystyrene tracer beads to the observation chamber and drove a single actuator with square-wave voltage pulses of ± 0.7 V at 5, 10, 20, 30, 40, and 50 Hz (**FIGURE 2**). High-speed movies were acquired at 330 fps and postprocessed to enhance contrast, followed by particle tracking using machine-learning based segmentation SAM2 (*37*) (more details in the *Supporting Information*). Within a fixed region of interest (ROI) directly above the actuator tip, three tracer particles with different distances from the actuator (P1–P3) were identified (**FIGURE 2A**, Error! Reference source not found.), and their *y*-positions tracked as a function of time, demonstrating that the particles follow the periodic bending of the actuator, with decreasing amplitude at larger particle-actuator distance (**FIGURE 2B, Movie S1**). We then probed different actuator frequencies and extracted the maximum (positive and negative) velocity *v* along the y-direction for each particle (**FIGURE 2C**). Additionally, we report the corresponding Stokes drag force *F* = 6*πηrv* acting on each particle of radius *r*, where *η* denotes the dynamic viscosity of water. This quantification reveals a maximum of the microactuator-generated flow at biologically relevant frequencies.

**FIGURE 2:**
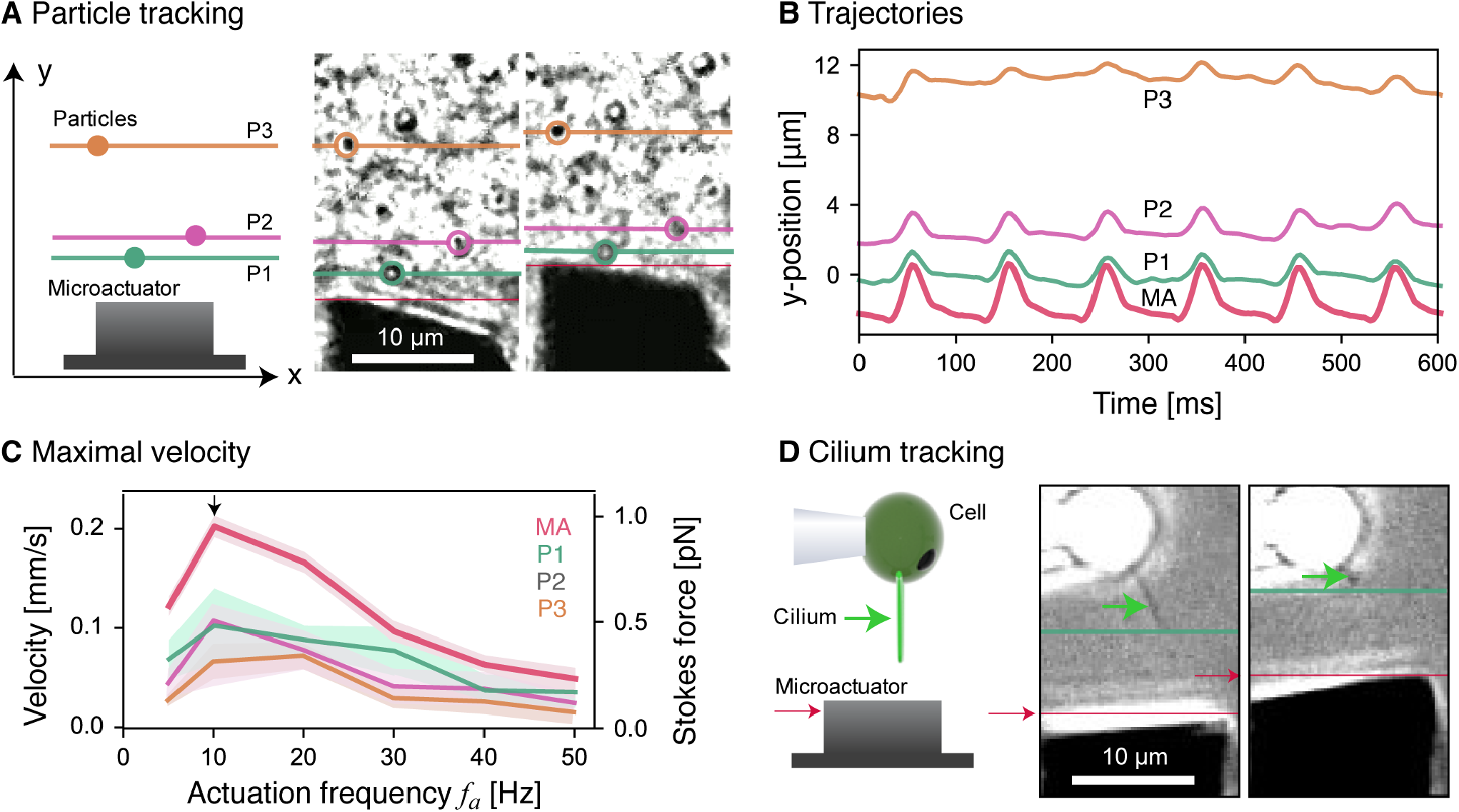
Quantification of tracer particles and cilium motion driven by nearby microactuation. **(A)** Schematic top view of the imaging geometry and two representative microscope images (at subsequent time points) showing 0.5-μm-diameter polystyrene tracer particles (P1-P3) within the region of interest (ROI) above the microactuator; colored lines indicate the tracked *y*-positions of P1–P3 used for subsequent analysis. **(B)** Time-resolved *y*-position of the microactuator tip (MA) and the three tracer particles (P1–P3) at a fixed driving frequency of 10 Hz. **(C)** Maximum velocities of the actuator and particles (left axis) and corresponding Stokes forces acting on the particles (right axis; mean±sd; shaded regions; *n*=66 cycles) as function of actuator frequency. **(D)** Schematic and two representative microscope images showing simultaneous tracking of a single cilium and the neighboring microactuator using the same ROI as in (B); colored lines indicate the tracked *y*-position of the cilium tip (green) and the microactuator tip (red).

Having established the microactuator-driven flow field, we investigated cilia responses to these local flows. For this, we used the same ROI to track the motion of a single cilium of a living *Chlamydomonas* cell positioned above an adjacent microactuator using a microcapillary (**FIGURE 2D**). To simultaneously segment and track both the cilium and the microactuator across all frames, we again used SAM2. Because SAM2 combines segmentation with instance-aware tracking, it can reliably follow deformable structures and maintain object identity even when the cilium transiently changes shape, or moves partially out of focus during the recording. To match the optimum of the microactuator-generated flow and the intrinsic cilia beat frequency we chose an outer-dynein-arm-deficient *Chlamydomonas reinhardtii mutant oda1* (from now on referred to as *Chlamydomonas* cell), whose intrinsic ciliary beat frequency is reduced to approximately 20–30 Hz compared to 50–60 Hz in wild type (*22,38*).

Having established the biohybrid experimental platform, we analyzed ciliary beating in the presence of microactuator generated flows for different driving frequencies in the range 15–25 Hz (with 0.5 Hz increments; for each voltage, 200 square-wave voltage pulses of ± 0.7 V were applied, with 2s pauses between runs (**Movie S2**)). **FIGURE 3** exemplifies the analysis for 20-Hz driving where the *y*-positions of the cilium tip and the microactuator tip are plotted as function of time (**FIGURE 3A**). Once the driving voltage is switched on, the electrochemical operating point of the microactuator slowly relaxes to a stable steady-state on a timescale of several seconds. This is reflected by a gradual change in the mean position of the microactuator (see also Error! Reference source not found.). For analysis, we define four time windows: ‘before actuation’ (gray), ‘transient regime’ (light-brown), ‘stable actuation regime’ (dark-brown), and ‘after actuation’ (blue). We automatically detected the minima of the cilium tip positions (green crosses) and actuator positions (red circles, see zoom-in in the lower panel of **FIGURE 3A)**.

**FIGURE 3:**
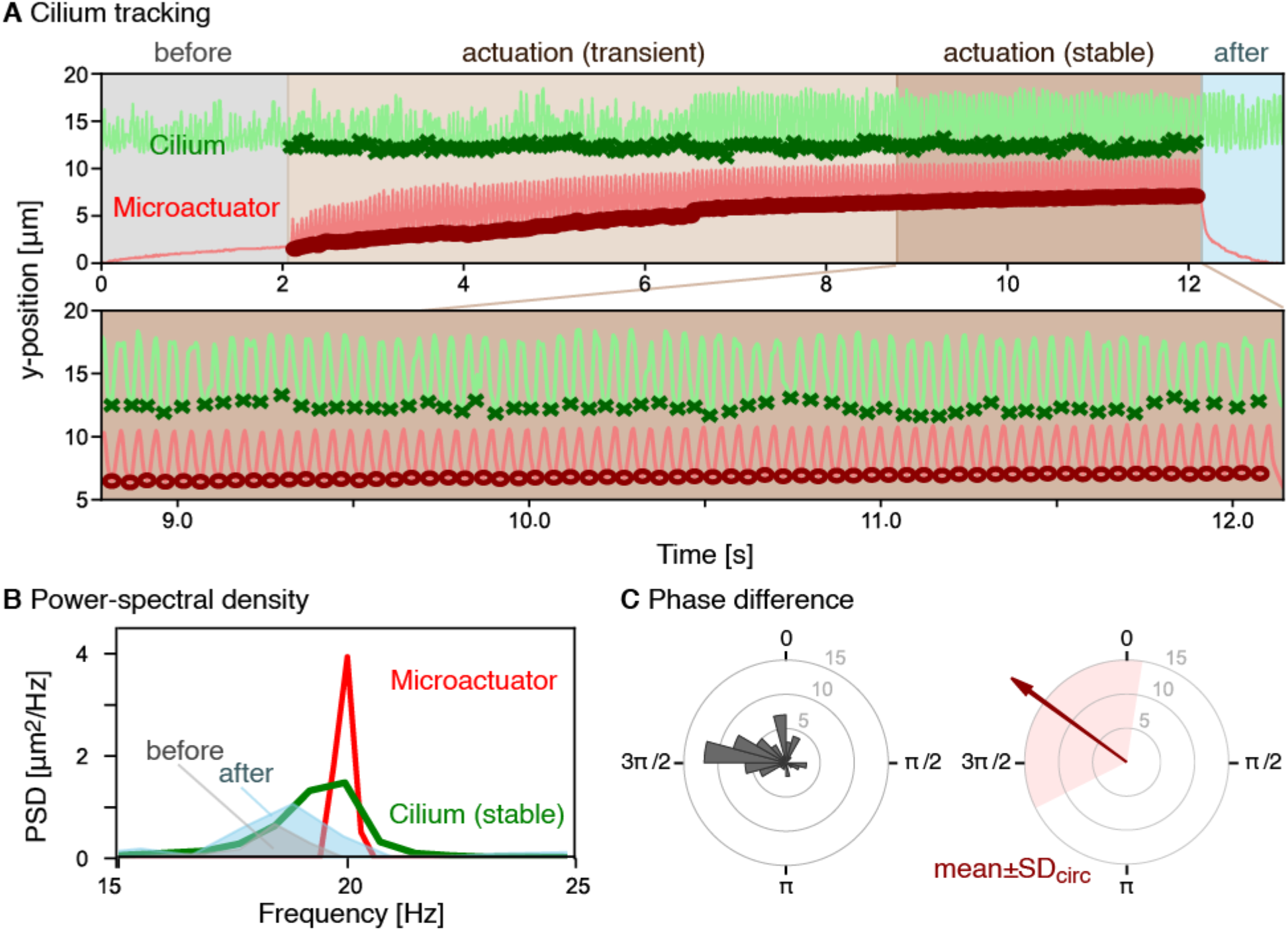
Hydrodynamic coupling between cilium and microactuator. **(A)** Time-resolved *y*-position of the cilium tip (green) and actuator tip (red) during a train of voltage pulses at a fixed 20-Hz driving frequency. Shaded regions denote (i) the pre-stimulus baseline (gray), (ii) the transient regime (light brown), (iii) the stable regime (dark brown), and (iv) the post-stimulus regime (blue). *Lower plot:* Zoom-in, displaying *y*-positions during the stable regime, with detected minima positions of the cilium (green crosses) and actuator (red circles) used to compute phase differences. **(B)** Power spectral densities (PSD) of cilium motion before and after stimulation (gray and blue, respectively) and in the stable regime (green), together with the PSD of the actuator in the stable regime (red). The ciliary peak shifts from its intrinsic frequency in the pre- and post-stimulus baseline toward the driving frequency of the microactuator, indicating frequency entrainment. **(C)** (left) Histogram of phase differences between cilium and microactuator (evaluated at minima of cilium tip *y*-position during the stable regime as marked in panel (A)). (right) Circular mean (red arrow, 1.68π) and circular standard deviation SD_circ_ (pink, 0.37π; circular variance 0.484) of the phase differences in graphical form.

### 2.1 Frequency entrainment

To characterize changes in cilium beat frequency upon actuation, we computed power-spectral densities (PSDs) of cilium and microactuator motion (**FIGURE 3B**). For cilium beating, PSDs were computed using Welch’s method (*39*) to reduce noise. For driving frequency of 20 Hz, we observe similar cilium PSDs before and after actuation. However, the cilium PSD during stable actuation (green, peak at approximately 19 Hz) is substantially different from the cilium PSDs before and after actuation but more similar to the PSD of the actuator (red). This suggests frequency entrainment to microactuator-generated oscillatory flow.

### 2.2 Phase-locking

To further study the entrainment between the microactuator and the cilium, we tested for phase-locking. For this, we define an oscillator phase ϕ(*t*) of the microactuator as follows: the minima of the *y*-position of the microactuator tip mark the completion of a full actuation cycle. At these minima, the phase ϕ(*t*) is set to subsequent integer multiples of 2π, and linearly interpolated in between (similar to the method described in (17)). We then compute the phase differences between the microactuator and the cilium by evaluating ϕ(*t*) whenever the *y*-position of the cilium attains a local minimum. The result of this “stroboscopic” analysis of phase differences for the stable regime is shown in **FIGURE 3C**. We additionally report the circular mean (1.68π) and the circular standard deviation SD_circ_ (0.37π) of these phase differences, which are computed using circular statistics. We repeated this analysis for the transient regime, during which the distance between microactuator and cilium is larger on average and obtained an increased circular standard deviation of 0.69π (**FIGURE S6)**. These findings are consistent with phase-locking of the beating cilium to the oscillatory flows generated by the microactuator during the stable regime, with a fixed, non-random phase relationship.

### 2.3 Frequency sweep

Our programmable microactuator allows us to generate custom driving frequencies. We tested cilium phase-locking for different driving frequencies *f*_actuator_ around the expected intrinsic cilium beat frequency *f*_cilium,0_ in the range of 15–25 Hz (with 0.5 Hz increments; 200 square-wave voltage pulses of ± 0.7 V for each frequency with 2-s pauses between actuation intervals). **FIGURE 4A** (left panel, red dots) shows the circular standard deviation (SD_circ_) of phase differences between the beating cilium and the microactuator as a function of *f*_actuator_ – *f*_cilium,0_. The analysis is analogous to Figure 3, with phase differences determined stroboscopically during the stable regime of actuation; *f*_cilium,0_ was determined as the mean of the beat frequencies determined from the PSDs of the beating cilium in the pauses between actuation intervals. Again, we find SD_circ_ indicative of phase-locking, with a tendency for smaller SD_circ_ if the microactuator frequency matches the intrinsic beat frequency of the cilium. As a negative control, virtually pairing timeseries data of equal duration of the same cilium without stimulation and of the microactuator during stable actuation, and conducting the same analysis again, resulted in consistently higher values of SD_circ_ (**FIGURE S6**).

**FIGURE 4:**
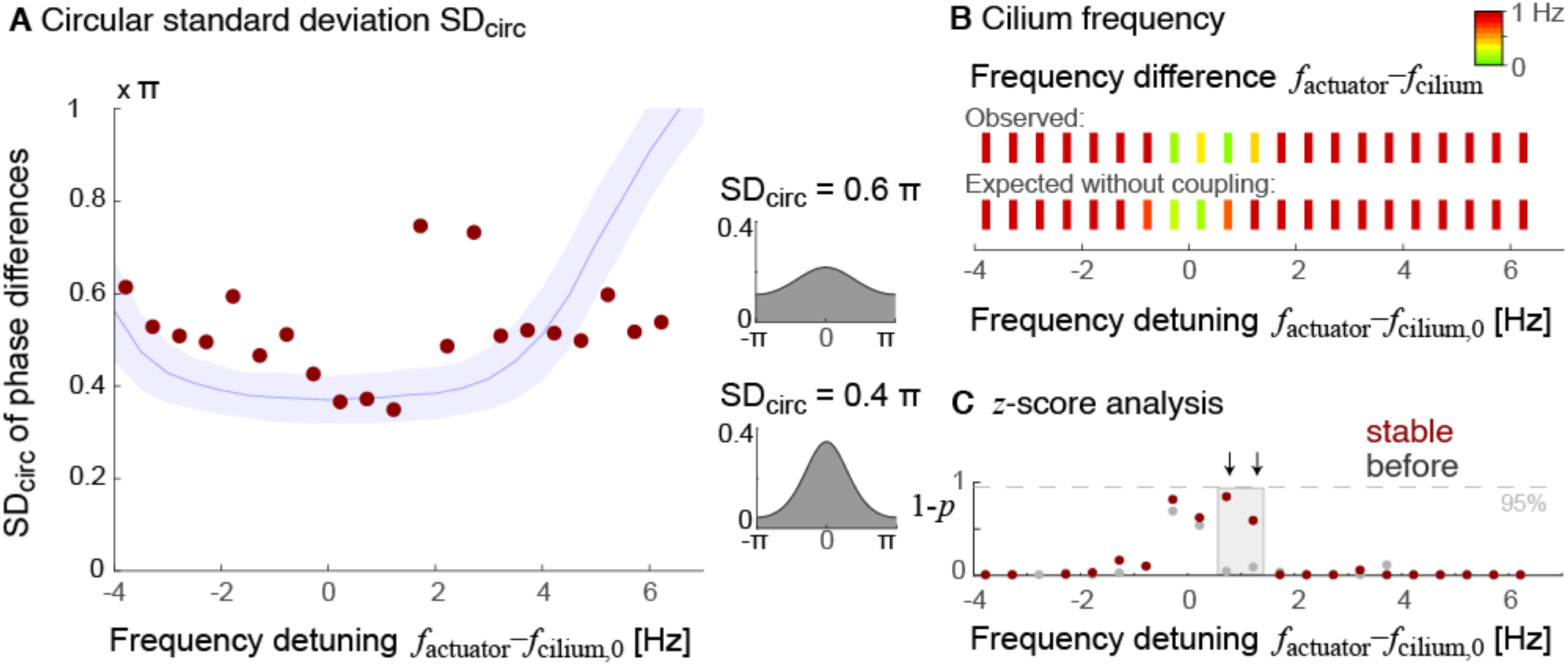
Frequency-dependent phase-locking between microactuator and cilium. **(A)** Circular standard deviation (SD_circ_) of phase differences between beating cilium and microactuator during the stable-actuation regime as a function of frequency detuning between the driving frequency *f*_actuator_ of the microactuator and the intrinsic beat frequency *f*_cilium,0_ of the cilium (red dots, *n*=66 cycles per condition). For reference, we additionally show simulation results from a minimal mathematical model of phase-locking (blue, mean±SD), see Eq. (1) (left panel). Representative von-Mises distributions for two values 0.4 and 0.6 of the SD_circ_ (right panel). **(B)** Observed frequency difference *f*_actuator_ – *f*_cilium_ between microactuator and cilium during the stable actuation regime (upper row, color coded) as well as expected difference if there was no coupling (lower row) as a function of frequency detuning *f*_actuator_ – *f*_cilium,0_. **(C)** Central probabilities 1–*p* corresponding to *z*-scores of normalized frequency differences as shown in panel (B), as function of frequency detuning *f*_actuator_ – *f*_cilium,0_, both for the regime of stable actuation (red) and before actuation (gray). Arrows point to the two detuning frequencies where the central probabilities differ most.

For further comparison, we include simulation results from a minimal mathematical model of phase-locking in the presence of noise (**FIGURE 4A**, right panel, blue curve), according to a modified Adler equation

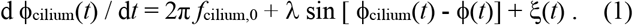

Here,ϕ_cilium_(*t*) denotes a simulatd time-resolved oscillator phase of the beating cilium, *f*_cilium,0_ is the intrinsic cilium beat frequency without actuation as above, ϕ(*t*) is the oscillator phase of the microactuator, which evolves according to d ϕ(*t*) / dt = 2π *f*_actuator_, λ is a generic coupling strength for the coupling between the microactuator-generated flows and the beating cilium, and ξ (*t*) is a Gaussian white noise term representing active phase fluctuations of the cilium beat (*22*), with < ξ (*t*) ξ (*t*’) > =2*D* δ (*t*-*t*’), *D* = 2π *f*_cilium,0_ / (2*Q*) with quality factor *Q* of cilium beating characterizing frequency jitter. The following parameters were used *f*_cilium,0_ = 18.73 Hz, λ = 0.3·2π *f*_cilium,0_ (same as the synchronization strength parameter determined in (*40*)), *Q* = 15 (smaller yet comparable to *Q* = 40 reported for *oda1* mutant in (*22*)); simulated time interval and number of data points matches experiment. Generally, a higher quality factor *Q* in the model results in lower SD_circ_, while higher coupling strength A results in both lower SD_circ_ and a wider trough of the U-shaped curve. Additionally, two reference distributions with prescribed values of SD_circ_ are shown (**FIGURE 4A**, left; von-Mises distributions (*39*)).

We additionally tested for frequency entrainment. **FIGURE 4B** shows the difference between the observed cilium beat frequency *f*_cilium_ during actuation in the stable regime, and the driving frequency *f*_actuator_ of the microactuator as a function of frequency detuning *f*_actuator_ – *f*_cilium,0_. For moderately small detuning **≲**1 Hz, the beating cilium adopted the driving frequency of the microactuator as its beat frequency. For larger detuning >1Hz, the microactuator-generated flows were not sufficient to impose the driving frequency on the beating cilium. To further quantify this observation, we computed *z*-scores, *z* = *f*_actuator_ – *f*_cilium_ / *s*, and central probabilities 1 – *p* (**FIGURE 4C**). Here, *p*(*z*) is the two-sided tail distribution of the standard normal distribution. The uncertainty *s* = 0.92 Hz of the estimated cilium beat frequency (without actuation, using observation times comparable to the duration of the stable actuation regime) was used to compute *z*-scores. We observed consistently higher values of 1 – *p* for the regime of stable actuation (red) than for the regime before actuation (gray) in the range of frequency detuning **≲**1 Hz corresponding to frequency entrainment. The observed range of frequency entrainment is reminiscent of an Arnold tongue (*41*), i.e., a window of frequency mismatches for which entrainment occurs, which may depend on the amplitude of driving. These results are typical for an analogous analysis for a second cell (see Error! Reference source not found.**Movie S3**). Similar Arnold tongues were previously reported for *Chlamydomonas* cilia (*13*) and cilia bundles (*42*) entrained by global oscillatory flows.

## 3 Discussion

We established and demonstrated a controllable, biocompatible microactuator capable of generating local oscillatory fluid flows, to probe the response of a single motile cilium to a local force. We characterized microactuator-generated flows by particle tracking and PIV, demonstrating that a single actuator produces measurable local oscillatory flow (with typical flow amplitudes of 50–100 μm/s at micron-distances from the actuator). Flow amplitudes exhibit a characteristic frequency dependence, which reflects the electrochemical/mechanical response of the microactuator, with a peak at biologically relevant frequencies of 10–20 Hz. The observed distance-dependence of microactuator-generated local flows can be used to tune the coupling strength in future experiments.

Placing motile cilia of living *Chlamydomonas* cells in the vicinity of a microactuator allowed us to probe cilia dynamics with and without actuation and at different driving frequencies. For small frequency detuning (small differences between the intrinsic beat frequency of the motile cilium and the driving frequency of the microactuator), we observed several characteristic signatures of frequency entrainment and phase-locking: (i) During actuation, the power-spectral density of ciliary dynamics aligned to that of the actuator, yet returned to its pre-actuation shape when the microactuator was switched off. This is indicative of reversible frequency entrainment. (ii) Analysis of phase differences between the beating cilium and the microactuator during stable actuation revealed a fixed phase-relationship between cilium and actuator, corroborating frequency entrainment. (iii) Frequency sweeping confirmed that frequency entrainment and phase-locking is most pronounced if the frequency detuning between cilium and the microactuator is small (on the order of 1 Hz). (iv) A minimal mathematical model of a noisy phase oscillator coupled to an external oscillatory driver (modeled using a modified Adler equation), reproduced the qualitative U-shaped dependence of the phase difference between cilium and microactuator as function of frequency detuning.

Relative to global oscillatory flows in channels, where entrainment is well established but geometry constrained (*13*), our platform enables (i) localized hydrodynamic forcing at cellular scales, (ii) rapid on/off switching without changing bulk flow conditions, and (iii) straightforward up-scaling to arrays of independently addressable microactuators through microfabrication. An additional contribution to the analysis of ciliary synchronization is our data analysis pipeline that combines machine learning-based tracking of shape-changing objects (here the biological cilium and the microactuator), reconstruction of an oscillator phase, and circular statistics. Automation of actuation and analysis allows for performing systematic parameter sweeps (frequency, waveform, distance, amplitude) and quantitative comparison across those conditions.

In conclusion, we have established a versatile experimental platform that enables controlled investigation of hydrodynamic synchronization at the cellular scale by combining wafer-scale, nanometer-thin Ti/Pt electrochemical microactuators with single-cell micromanipulation and high-speed imaging. This approach bridges the current gap between studies using either global-flow entrainment (*13*) or external global magnetic fields (*26*) and local, biologically driven flow, while overcoming key limitations of each. In particular, the ability to generate hydrodynamic forcing *in situ*, with electronically programmable frequency and amplitude and under biocompatible sub-Volt operation, enables systematic exploration of synchronization phenomena in well-controlled environments that was previously inaccessible.

Looking forward, the microactuators presented here offer a clear pathway toward actuator arrays with individually addressable control, enabling engineered metachronal pumping in microfluidic systems and causal studies of synchronization in larger ciliary collectives. Beyond cilia, such platforms could be used to apply precisely defined hydrodynamic stimuli to a wide range of biological cells, facilitating studies of mechanotransduction and cell-level responses. The development of multiplexed, programmable actuation schemes will further allow the generation of complex spatiotemporal flow patterns, advancing both fundamental research and applications in microfluidics, diagnostics, and biohybrid systems.

## 4 Materials and Methods

### 4.1 Substrate treatment

Square borosilicate glass plates (50 mm × 50 mm, 0.50 mm thickness) were used as substrates. Slides were rinsed with isopropyl alcohol (IPA, technical grade), thoroughly flushed with deionized water, and dried under a nitrogen stream. Immediately thereafter, an adhesion-promoter (TI PRIME, MicroChemicals) was dispensed onto the substrates and spin-coated (e.g., 3000 rpm, 30 s). The coated substrates were soft-baked on a leveled hotplate at 120 °C for 2 min to remove residual solvent and activate the coupling chemistry. TI PRIME forms an ultrathin coupling layer improving subsequent resist adhesion. Substrates were allowed to cool to room temperature in the cleanroom.

### 4.2 Microactuators fabrication and release

The microactuators were fabricated using microfabrication technology in the clean-room facilities of IFW Dresden in a fully monolithic, wafer-scale process. TI Prime–treated glass substrates were first spin-coated with a positive photoresist (AZ5214E, Microchemicals GmbH, Ulm, Germany) at 4500 rpm for 45 s and prebaked at 90 °C for 5 min. The resist was then patterned by maskless UV lithography using an MLA101 (Heidelberg Instruments) and developed in MIF 726 developer, followed by a DI-water rinse and N_2_ drying. Subsequently, a 200 nm aluminium layer was deposited by DC magnetron sputtering (NanoPVD) and structured by lift-off in acetone, followed by rinsing in isopropanol (IPA) and drying with nitrogen. To definine the Ti/Pt microactuators, a second lithography step with AZ5214E (positive resist; 90 °C, 5 min prebake; MLA101 exposure; MIF 726 development) was performed on top of the Al-patterned substrates. An 18 nanometer-thin Ti/Pt bilayer was then deposited by DC magnetron sputtering (NanoPVD) and patterned by lift-off in acetone with an IPA rinse, yielding the final Ti/Pt microactuator structures. Each glass wafer yielded 18 chips, each containing hundreds of microactuators. The wafer was scribed and diced with a glass cutter, and the individual chips were immersed in MIF 726 developer to remove the aluminum sacrificial layer, thereby triggering the self-assembly of the microactuators. After release, the chips were thoroughly rinsed in DI water and stored in 99% ethanol until use in subsequent experiments in the 3D-printed chamber.

### 4.3 Chamber fabrication

3D-printed double chambers were fabricated by the Microstructure Facility, a core facility of the CMCB Technology Platform at TU Dresden. The structures were modeled using BioCAD software (RegenHU) and printed with SE 1700 silicone (Dow Dowsil™) on a RegenHU 3D bioprinter. Printing was performed directly onto 60 × 24 mm coverslips, which had been pre-cleaned in 1 % Helmanex solution and washed in distilled water using an ultrasonic bath. First, an inner wall with a height of 400 μm was printed to form two separate reservoirs within the chamber. This was followed by printing the outer frame with a total height of 4 mm. A conical nozzle with an inner diameter of 400 um was used to print the structures. The printed chambers were cured at 60°C for 4 hours and at 78 °C for 2 hours. For experiments, released microactuator chips were placed face-up on the chamber floor and positioned between two screws that had been glued to the bottom of the chamber. A 3D-printed holder carrying a 0.5 mm Pt wire (working electrode) and featuring two through-holes was then placed onto the screws, and nuts/bolts were tightened to press the Pt wire onto the actuator contact area, providing a stable electrical connection inside the fluid.

### 4.4 Electrical control of microactuators

Actuation of the Ti/Pt microactuators were controlled with an Agilent B2902A source-measure unit (SMU) using a custom Python program (PyVISA-based instrument control with a Tkinter GUI for parameter entry, live plotting, and automated data logging). At program start, the SMU was identified and initialized via standard SCPI commands (instrument query, reset, and clear), and the output channel was configured for 4-wire (remote-sense) operation. All experiments were conducted in a three-electrode configuration: the Pt actuator contact served as the working electrode (WE), a platinum 0.5 mm wire as the counter electrode (CE), and a pseudo Ag/AgCl reference electrode (RE) provided a stable reference potential. Before applying any waveform, the program first performed an open-circuit potential (OCP) check in which the SMU sources 0 A (current-source mode) while measuring the resulting WE potential versus time. This step was used as a quick verification of correct electrical contact and a stable electrochemical connection in the chamber. To “activate” the actuator response and reproducibly cycle the material between its reduced and oxidized states, the program then executed cyclic voltammetry (CV) for typically ≥20 cycles. In the implementation, the CV waveform was generated as a discretized voltage list defined by the start, upper, lower, and stop potentials with a defined step size and scan rate. After activation, the microactuators were driven using square-wave pulsing between a high (oxidizing) and low (reducing) potential to produce large mechanical amplitudes, especially at higher drive frequencies. In the code this is implemented via the SMU’s built-in pulse shape mode, where the user specifies the top and bottom voltages, frequency, duty cycle, number of cycles, and current compliance. The pulse timing (period and width) is derived from the chosen frequency and duty cycle, and the SMU is triggered by an internal timer to generate a stable, repeatable pulse train. During pulsing, the program acquires voltage, current, and time stamps for each pulse and saves the full dataset automatically.

### 4.5 Imaging

The microactuators, particles, and *Chlamydomonas* cells were imaged using phase-contrast microscopy set up on an inverted Nikon ECLIPSE Ti2-E microscope, using a Nikon S Plan Fluor ELBW 60x, 0.7 NA lens in combination with a 1.5x or 1.0x tube lens and a Nikon LWD 0.52 air condenser. High speed movies were recorded using a pco.dimax CS4 high-speed camera, overview movies of the actuators were recorded with a pco.edge 4.2 camera. In both cases, the sample was illuminated using a Sola SE2 light engine (Lumencor) combined with a 496 LP filter (Semrock, Brightline) and hardware was controlled by NIS-Elements.

### 4.6 Tracer particle experiments

To quantify actuator-generated oscillatory flows, the chamber medium was seeded with monodisperse polystyrene tracer beads (0.5 μm diameter, Sigma-Aldrich, 95585-5ML-F). For each experiment, 10 μL of the particle stock suspension was diluted into 3 mL of fresh TAP medium directly in the chamber. Highspeed videos (330 fps) were recorded while driving the actuator at the desired frequency. Tracer trajectories were extracted using a SAM2 machine learning pipeline. For each frequency, particle displacement traces were segmented into positive and negative half-cycles by detecting maxima and minima, and peak velocities were computed separately for the positive and negative directions. Three representative particles at distinct actuator distances were selected for reporting; particles were ranked by their mean distance to the microactuator (computed from the average of the maxima- and minima-based measured distances). For each tracked particle, the mean ± standard deviation of the positive and negative peak velocities was calculated across the recording. To relate measured velocities to an effective hydrodynamic forcing scale, velocities were converted to a Stokes drag force.

### 4.7 Cell growth, capture and positioning

For holding the cells, a microfluidic glass micropipette (MGM 1A, FIVE photon Biochemicals), with a tip-length of 25μm and an opening diameter of 3μm was mounted on a Narishige, Coarse Manipulator (MN-4), which was mounted to a Narishige three-axis oil hydraulic Micromanipulator (MM)-303). The suction force was generated using a micro-injector (Eppendorf CellTram-oil). We used the *oda1* mutant of *Chlamydomonas reinhardtii* (strain CC-2228 mt- (*43*), obtained from chlamycollection.org) for all experiments. Cells were grown in a 300 ml Tris-Acetate-Phosphate (TAP).

medium under a 12/ 12 hour light/dark cycle and illuminated by an LED lightpad (Light Therapy Lamp, 10,000 LUX) under constant air-bubbling at 23°C, to a final density of approximately 10^6^ cells/ml (*36*).

For each experiment, a 15 μL aliquot of the culture was taken and added to 3 mL of fresh TAP medium in the 3D-printed chamber containing the microactuator. Individual cells were then captured at the tip of a glass microcapillary by applying gentle suction with the CellTram microinjector. The microcapillary holding the *Chlamydomonas* cell was then positioned with the three-axis micromanipulator to bring the cell as close as possible to the microactuator while maintaining a small gap, such that even at maximal ciliary extension the cilia did not contact the microactuator. Using the custom python program, an automatic train of electrical pulses (from the SMU) starting from lower frequencies to higher frequencies with predefined number of cycles and stall times in between were applied on the microactuator and one 230 fps movie was captured for all of the frequencies.

### 4.8 Segmentation using SAM2

The experiment videos of microactuators, tracer-particles, and cilia were segmented and tracked using SAM2 (Segment Anything Model 2), a deep-learning video segmentation model that generates object masks from sparse user prompts (e.g., clicks on the object) and then propagates these masks across frames. SAM2 was installed on a Linux workstation with GPU acceleration (CUDA), and used through a custom, locally hosted GUI written in Python. A custom Gradio-based GUI provided an end-to-end workflow, namely (i) importing a recorded video, (ii) extracting the frames using FFmpeg into an image sequence, (iii) choosing an annotation start frame and an end frame to define the tracking segment, and (iv) compute SAM2 features for that frame range (model state initialization). Objects were defined interactively by placing positive and negative point prompts on the annotation frame with live mask preview; additional objects (e.g., cilium, microactuator, and selected tracer particles) were added sequentially as separate object IDs. After annotation, SAM2 was run in video-prediction mode to generate per-frame masks for each defined object. The application automatically saved binary mask PNGs for each object and frame (organized into dedicated subfolders for cilia and actuator), and also generated an overlay “tracked video” (masks overlaid on the raw frames).

Before image segmentation, we pre-processed microscope videos to enhance bead contrast and suppress high-frequency noise, using contrast-limited adaptive histogram equalization (CLAHE) with an optimized clip limit followed by a median filter.

### 4.9 Computation of power-spectral densities

Power-spectral densities (PSD) of the microactuator were computed as single periodogram/FFT of the tracked *y*-position of the microactuator tip. PSDs of beating cilia were computed form the tracked *y*-position of the cilium tip using Welch’s method (*27*), which accounts for their frequency jitter. Welch’s method splits the signal into overlapping, windowed segments and averages their periodograms to reduce the variance of the spectral estimate and suppress noise in the signal. The resulting PSDs were then smoothed using a Savitzky–Golay filter and the dominant peaks fitted with Lorentzian functions. The peak positions of these Lorentzian fits define the dominant frequency of microactuator and cilium in the different actuation regimes.

### 4.9 Statistical methods

For each analyzed time window (e.g., transient and stable actuation regime), we quantified how tightly the cilium phase followed the microactuator drive by computing the distribution of phase differences and summarizing it using circular statistics. Phase differences were extracted from the tracked *y*(*t*) positions using an event-based definition that is robust to waveform distortion and amplitude modulation. First, local minima were detected using peak finding in the time series of *y*(*t*) positions of the microactuator and the cilium, respectively. The corresponding timepoints of these minima mark the completion of a full oscillation cycle and were assigned an unwrapped event phase

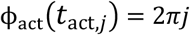

for the microactuator, and

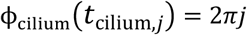

for the cilium, where *t*_act,*j*_ and *t*_cilium,*j*_ are the timepoints of subsequent minima. To compute relative phase differences, the microactuator event phase was linearly interpolated to the times of cilium minima, ϕ_act_(*t*_cilium,*j*_), and the phase difference computed as:

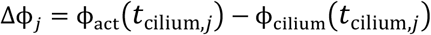

Given *n* phase differences Δϕ_*j*_, we computed the circular standard deviation using the standard formula

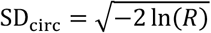

and the circular variance as 1-*R*, where 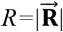 denotes the length of the mean resultant vector

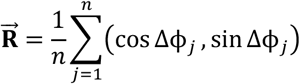

Here, *R*=1 and SD_circ_=0 indicates perfect phase-locking (all Δϕ_j_ identical), *R*<1 and SD_circ_>0 indicates a broader, less phase-locked distribution.

## 5 Acknowledgment

We thank Tomasz M. Basiewicz (Microstructure Facility, CMCB Technology Platform, TU Dresden) for 3D-printing the experimental chambers. We further acknowledge the support of the Research Technology Department of the Leibniz Institute for Solid State and Materials Research (Leibniz IFW) Dresden and the IFW cleanroom team led by Ronny Engelhard for their assistance in developing the experimental setup. B.M.F. was supported by the Deutsche Forschungsgemeinschaft (DFG, German Research Foundation) under Germany’s Excellence Strategy-EXC-2068-390729961 (Cluster of Excellence “Physics of Life”, TU Dresden), and through a DFG Heisenberg grant (421143374). M.M.-S. acknowledges financial support from the European Union’s Horizon 2020 research and innovation program (ERC Starting Grant Nr. 853609), the HORIZON-MSCA-2022-COFUND-101126600-SmartBRAIN3 and DRESDEN-concept.

