## Supporting Information for "Programmable microactuators phase-lock cilia to local oscillatory flow"

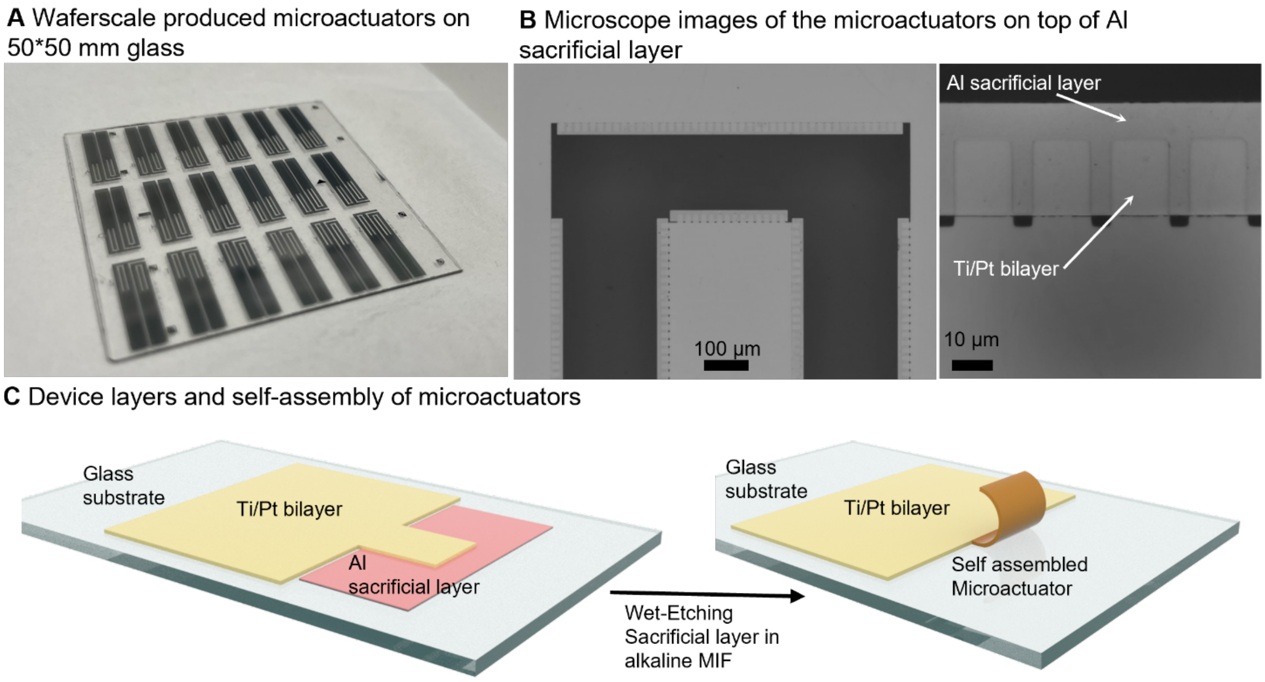


**FIGURE S1.** Wafer-scale fabrication and release of ultrathin Ti/Pt microactuators on glass. (A) Photograph of a microactuator array fabricated on a 50 × 50 mm glass substrate. (B) Representative bright-field microscope overview (left) and magnified view (right) of patterned Ti/Pt bilayers on top of an Al sacrificial layer prior to release; arrows indicate the Al sacrificial layer and Ti/Pt bilayer. (C) Schematic of the device stack (glass / Al sacrificial layer / Ti/Pt bilayer) and release process: wet etching of the Al sacrificial layer in alkaline metal-ion–free developer (MIF) triggers self-assembly of the freestanding microactuator.

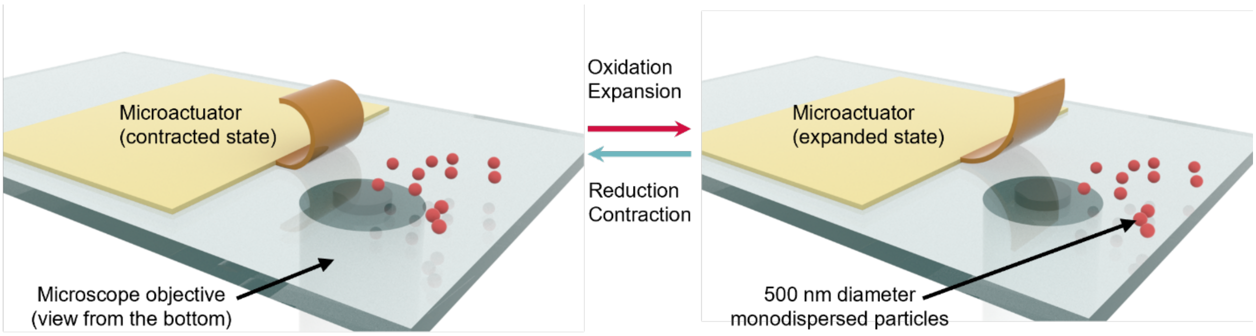


**FIGURE S2.** 3D schematic of tracer-particle tracking around a microactuator. The Ti/Pt microactuator switches reversibly between a contracted state (left) and an expanded state (right) through oxidation-driven expansion and reduction-driven contraction, generating local oscillatory flow in the surrounding medium. Monodisperse tracer particles (500 nm diameter) are imaged from below through the glass substrate with an inverted microscope objective, and their trajectories are used to quantify microactuator-induced fluid motion in the vicinity of the actuator tip.

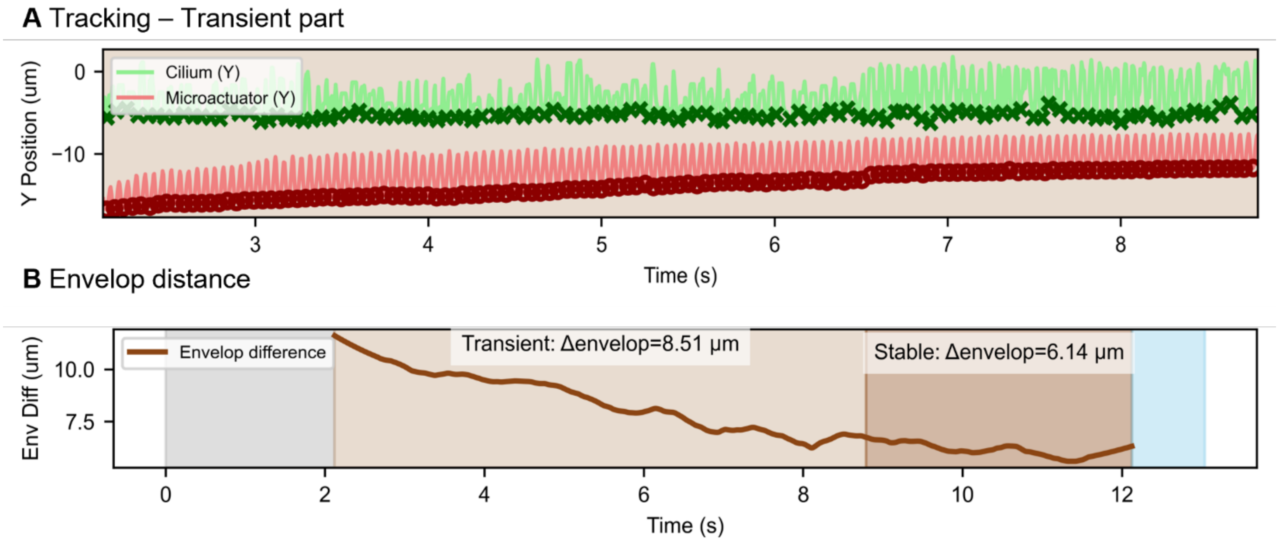


**FIGURE S3.** Identification of transient and stable response regimes from the envelope separation of cilium and microactuator. (A) Time-resolved y-position of the cilium tip (green) and microactuator tip (red) during periodic actuation, highlighting an initial transient state in which the distance decreases to finally reach a quasi-steady stable state. (B) Difference between the lower signal envelopes (envelope distance) as a function of time, used to quantify convergence from the transient regime (light brown) to a stable regime (dark brown); gray and blue shaded regions indicate the pre- and post-actuation intervals, respectively. Reported values give the mean envelope distance in the transient and stable windows.

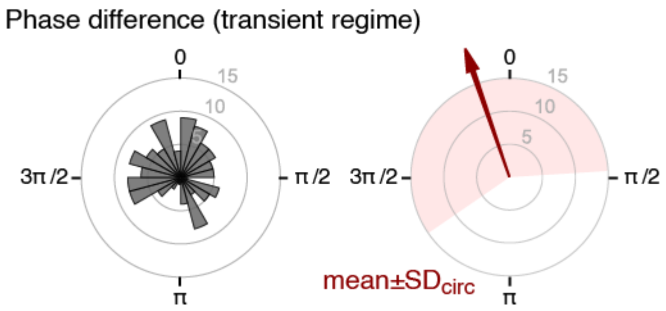


**FIGURE S4.** Histogram of phase differences between cilium and microactuator as shown in Fig. 3C, yet for the transient regime of actuation. The second plot (red arrow, 1.89π) and circular standard deviation $SD_{circ}$ (pink, 0.59π; circular variance 0.816) of these phase differences in graphical form.

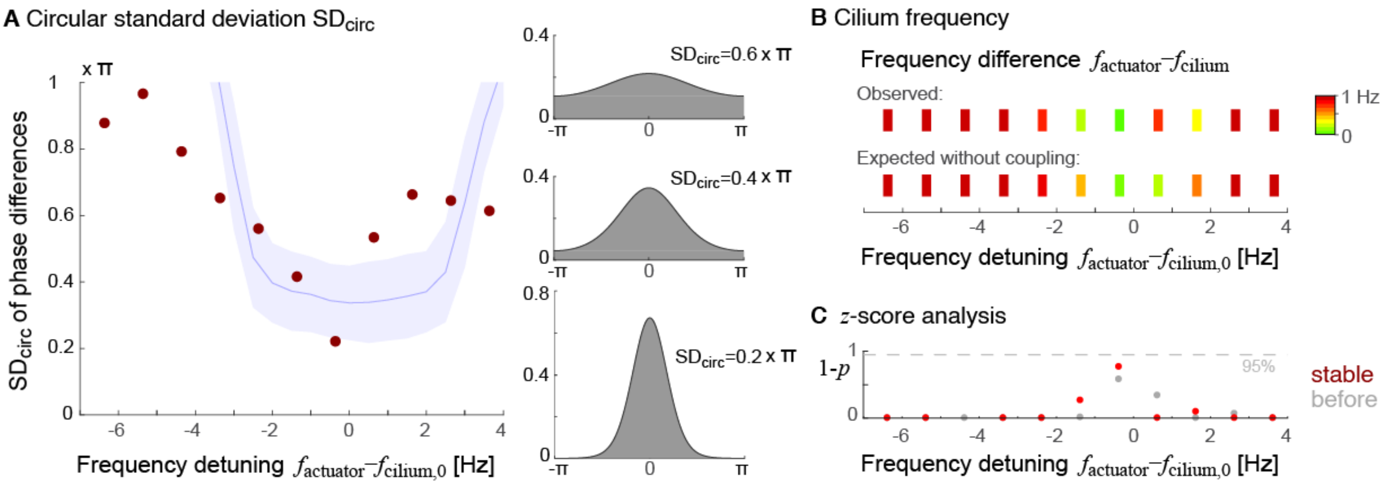


**FIGURE S5.** Frequency-dependent entrainment and phase-locking of a second *Chlamydomonas* cilium to microactuator forcing drive frequencies 10 to 20 Hz with 1Hz steps with the intrinsic frequency of cilium around 16 Hz, analogous to Fig. 4. **(A)** Circular standard deviation ($SD_{circ}$) of phase differences between beating cilium and microactuator during the stable-actuation regime as a function of frequency detuning between the driving frequency $f_{actuator}$ of the microactuator and the intrinsic beat frequency $f_{cilium,0}$ of the cilium (red dots, $n$=66 cycles per condition). For reference, we additionally show simulation results from a minimal mathematical model of phase-locking (blue, mean±SD), see Eq. (1) (left panel). Representative von-Mises distributions for three values $0.2\pi$ , $0.4\pi$ and $0.6\pi$ of the $SD_{circ}$ (right panel). **(B)** Observed frequency difference $f_{actuator} - f_{cilium}$ between microactuator and cilium during the stable actuation regime (upper row, color coded) as well as expected difference if there was no coupling (lower row) as a function of frequency detuning $f_{actuator} - f_{cilium,0}$. **(C)** Central probabilities $1-p$ corresponding to $z$-scores of normalized frequency differences as shown in panel (B), as function of frequency detuning $f_{actuator} - f_{cilium,0}$, both for the regime of stable actuation (red) and before actuation (gray).

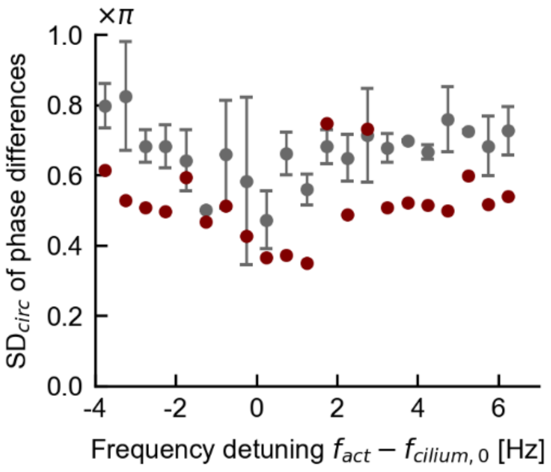


**FIGURE S6.** Randomized control for Figure 4. Circular standard deviation $SD_{circ}$ of cilium–actuator phase differences in the stable regime versus frequency detuning identical to Figure 4A (red), as well as random control (gray) calculated analogously but using cilium–actuator phase differences obtained by pairing time series of an unstimulated cilium and a time series of microactuator position during stable actuation; mean±sd, $n$=3 time series.

**Text for Fig. S6:** To provide a negative control (no coupling between actuator and cilium) that preserves finite-time effects, we constructed uncoupled surrogate datasets from an unstimulated recording of the same cilium (no voltage applied to the microactuator). The unstimulated trace (~12 s) was split into three segments with duration matched to the longest stable actuation window in the frequency sweep for the case of actuation at 15 Hz ($T_{\text{stable-segment}} \approx 4.7$ s), thereby controlling for the change in phase dispersion expected from limited observation time and intrinsic phase noise. For each driving frequency, each unstimulated cilium segment was then analyzed together with a same-length microactuator segment from the corresponding stable time window, using the microactuator minima only to define the reference phase timebase $\phi_{act}(t)$ for the stroboscopic evaluation. Because the cilium segment was recorded without actuation, this pairing contains no physical coupling, and it merely reproduces the identical analysis pipeline and time-windowing used for the actuated data. We report the mean ± SD of $SD_{circ}$ across the three surrogate segments as the control in **Figure S6** (left panel, gray dots).
